# An epigenetic mechanism for cavefish eye degeneration

**DOI:** 10.1101/199018

**Authors:** Aniket V. Gore, Kelly A. Tomins, James Iben, Li Ma, Daniel Castranova, Andrew Davis, Amy Parkhurst, William R. Jeffery, Brant M. Weinstein

## Abstract

Coding and non-coding mutations in DNA contribute significantly to phenotypic variability during evolution. However, less is known about the role of epigenetics in this process. Although previous studies have identified eye development genes associated with the loss of eyes phenotype in the Pachón blind cave morph of the Mexican tetra *Astyanax mexicanus*^1-6^, no inactivating mutations have been found in any of these genes^2,3,7-10^. Here we show that excess DNA methylation-based epigenetic silencing promotes eye degeneration in blind cave *Astyanax mexicanus.* By performing parallel analyses in *Astyanax mexicanus* cave and surface morphs and in the zebrafish *Danio rerio,* we have discovered that DNA methylation mediates eye-specific gene repression and globally regulates early eye development. The most significantly hypermethylated and down-regulated genes in the cave morph are also linked to human eye disorders, suggesting the function of these genes is conserved across the vertebrates. Our results show that changes in DNA methylation-based gene repression can serve as an important molecular mechanism generating phenotypic diversity during development and evolution.

---

Subterranean animals offer an excellent opportunity to study morphological, molecular and physiological changes that allow organisms to adapt to unique environments. Loss of eyes is one of the most common morphological features of cave-adapted animals, including many fish species. Blind cave fish (CF) morphs of *Astyanax mexicanus* evolved from surface fish (SF) during a few million years of isolation in dark Mexican caves^11^, with recent studies suggesting that regression of eyes evolved as part of a strategy to conserve energy in fish adapted to dark and nutrient deficient caves^12^. Although a number of studies have examined molecular mechanisms underlying eye loss in Pachón cave-derived *Astyanax mexicanus* CF, recent sequencing of the Pachón CF genome and other studies revealed no inactivating null mutations in essential eye development genes^2,3,7-9^. In contrast, genome sequencing of another subterranean animal, the naked mole rat *Heterocephalus glaber,* showed combined functional loss of more than a dozen key eye genes due to inactivating mutations^13^. These findings suggest the possibility that epigenetic rather than genetic changes may mediate eye loss in Pachón cave fish. To test this possibility, we used CF and SF morphs of *Astyanax mexicanus* as well as wild type and DNA methylation and demethylation-deficient zebrafish *Danio rerio* to examine whether DNA methylation regulates eye formation, and whether eye loss in Pachón cave fish evolved at least in part through hypermethylation of key eye genes.

At 36 hpf *Astyanax mexicanus* CF and SF embryos are superficially indistinguishable with properly formed lenses and optic cups in both morphs **(Fig. 1a,b)**. By five days of development, however, degeneration of eye tissue is clearly evident **(Fig. 1c,d)**, and by adulthood CF eyes are completely absent **(Fig. 1e,f)**^14^. Eye regression is preceded by decreased expression of a number of different eye-specific genes, including the crystallins *crybb1, crybb1c,* and *cryaa* (**Fig. 1g**). The expression of large sets of genes can be repressed by DNA methylation based epigenetic silencing, as perhaps most famously shown for X chromosome inactivation^15^. New epigenetic DNA methyl “marks” are added by specific enzymes known as *de novo* DNA methyltransferases (DNMTs)^16^, and we recently showed that one of these enzymes, *dnmt3bb.1,* is expressed in zebrafish hematopoietic stem and progenitor cells (HSPC) where its loss leads to failure to maintain HSPCs^17^. We found that *dnmt3bb.1* is also expressed in the ciliary marginal zone (CMZ) of the developing eye, a specialized stem cell-containing tissue surrounding the lens that is responsible for generating neurons and other eye cell types^18-20^ **(Fig. 1h)**. In addition to their hematopoietic defects *dnmt3bb.1*^*y258*^ null mutant larvae and adults have enlarged eyes compared to their wild type (WT) siblings **(Fig. 1i-k)** with retinal hyperplasia **(Extended Data Fig. 1),** and increased expression of a number of different eye genes, including *opn1lw1, gnb3a,* and *crx* **(Fig. 1l)**. Interestingly, the closely related *Astyanax mexicanus dnmt3bb.1* gene shows 1.5-fold increased expression in Pachón CF compared to SF **(Fig. 1m)**. The inverse correlation between eye size and *dnmt3bb.1* expression in *Danio rerio* and *Astyanax mexicanus* led us to hypothesize that excessive Dnmt3bb.1-dependent methylation may globally repress expression of eye genes in CF.

**Figure 1.**
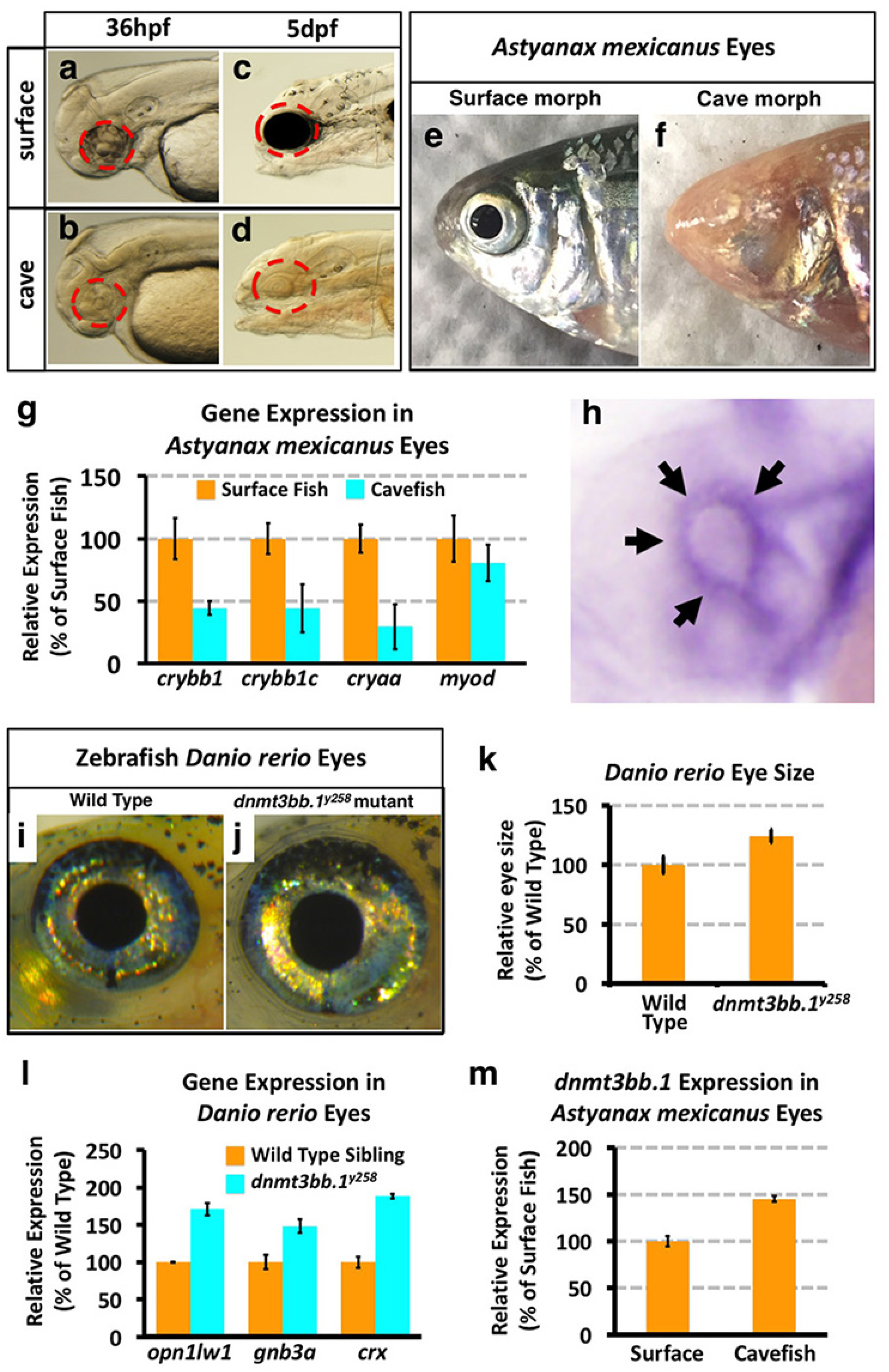
Eye phenotypes in *Astyanax mexicanus* surface and cave fish, and in zebrafish *Danio rerio* wild type and *dnmt3bb.1*^*y258*^ mutant animals. **a-d,** Transmitted light photomicrographs of the heads of 36 hpf (a,b) and 5 dpf (c,d) surface (a,c) and cavefish (b,d) morphs of *A. mexicanus.* Dotted red circle mark the developing eyes in CF and SF. **e,f,** Photographic images of the heads of adult surface (e, with eyes) and cave (f, eyeless) morphs of *A. mexicanus.* **g,** Quantitative RT-PCR analysis of the percent relative expression of *crybb1, crybb1c, cryaa,* and *myod* in isolated heads from 54 hpf surface (orange columns) and cave (blue columns) morphs of *A. mexicanus,* normalized to surface fish levels. **h,** Whole mount *in situ* hybridization of a 36 hpf zebrafish eye probed for *dnmt3bb.1,* showing expression in the ciliary marginal zone (CMZ, arrows). **i,j,** Photographic images of the eyes of adult wild type sibling (i) and *dnmt3bb.1*^*y258*^ mutant (j) zebrafish. **k,** Quantitation of eye size in three week old wild type sibling and *dnmt3bb.1*^*y258*^ mutant animals. **l,** Quantitative RT-PCR analysis of the percent relative expression of *opn1lw1, gnb3a,* and *crx* in adult wild type sibling (orange columns) and *dnmt3bb.1*^*y258*^ mutant (blue columns) zebrafish eyes, normalized to wild type sibling levels. **m,** Quantitative RT-PCR analysis of the percent relative expression of *dnmt3bb.1* in surface and cave morphs of *A. mexicanus,* normalized to surface fish levels. All images are lateral views, rostral to the left.

To more comprehensively examine whether increased DNA methylation correlates with reduced eye gene expression in CF versus SF, we performed combined RNAseq and whole genome bisulfite sequencing on RNA and DNA simultaneously co-isolated from 54 hpf CF or SF eyes, during a critical period for eye development in Pachón CF^14^ **(Fig. 2a)**. RNAseq analysis confirmed increased expression of *dnmt3bb.1* in CF eyes **(Fig. 2b,c)**. The RNAseq data also revealed that a large number of different eye development genes show reduced expression in CF eyes **(Fig. 2c and Supplementary data 1)**. As in the naked mole rat, visual perception (GO:0007601) and electrophysiology of eye are the top down-regulated biological processes found using Gene Ontology (GO) **(Fig. 2d)** and Ingenuity Pathway Analysis **(Extended Data Fig. 2)**, respectively. Ingenuity pathway analysis also predicted that the phototransduction pathway is the most affected signaling pathway in cavefish eyes compared to surface eyes **(Extended Data Fig. 2).**

**Figure 2.**
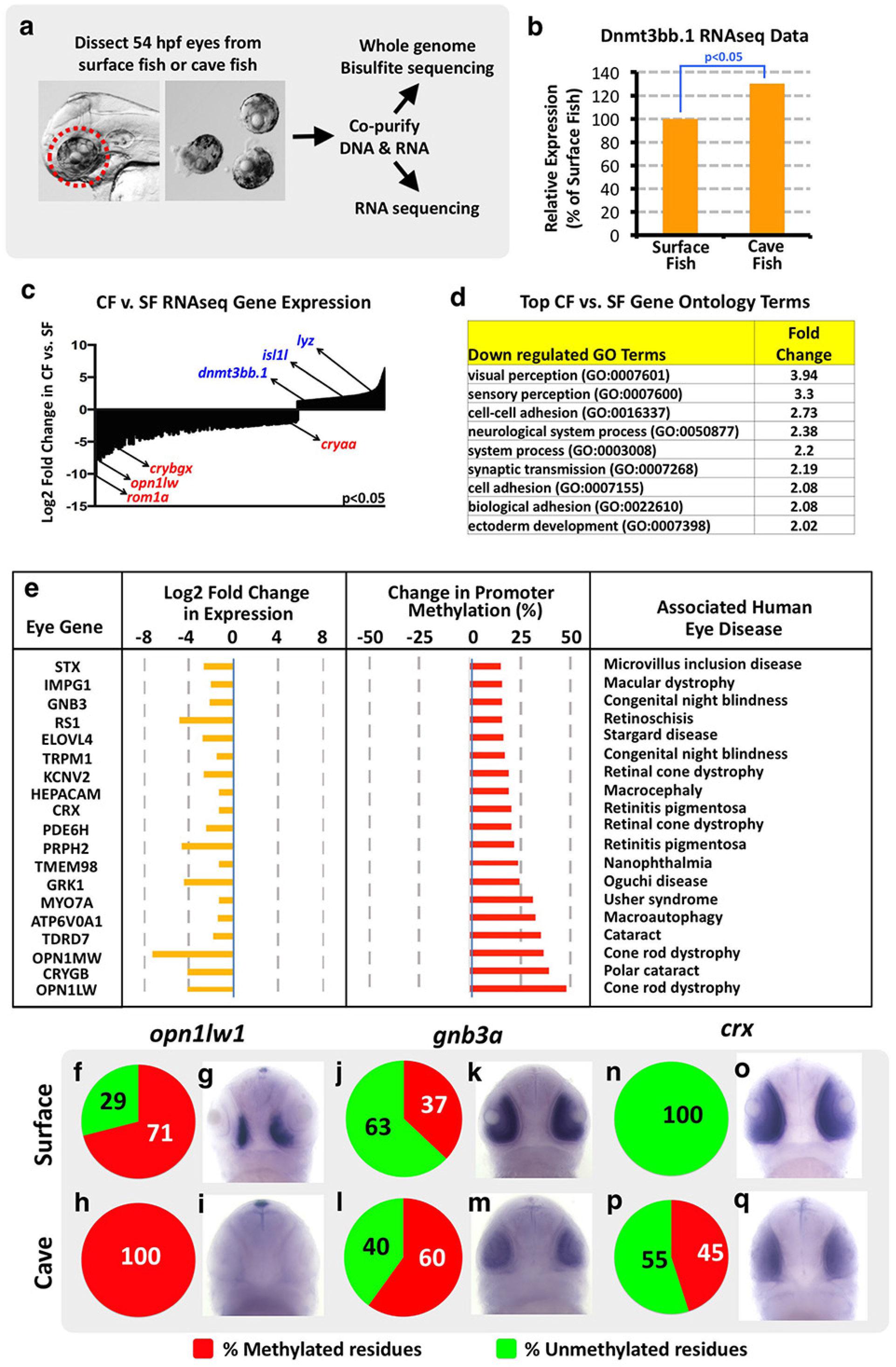
Gene expression changes in cave versus surface fish morphs of *Astyanax mexicanus.* **a,** Diagram showing the workflow for obtaining larval eyes from *A. mexicanus* and for co-isolating eye DNA and RNA for whole genome assessment of DNA methylation and gene expression, respectively. **b,** Percent relative expression of Dnmt3 in the eyes of *A. mexicanus* cave vs. surface fish morphs by comparison of their respective RNAseq data sets, normalized to surface fish levels. **c,** Log2 fold differential expression (p<0.05) of genes in *A. mexicanus* cave vs. surface fish morph RNAseq data sets, with down-regulated (red) or up-regulated (blue) expression of selected genes in CF noted. **d,** Listing of the Gene Ontology (GO) terms showing the greatest down-regulation in cave fish compared to surface fish. **e,** Nineteen genes with both substantially increased methylation within the 2 Kb of genomic DNA upstream from the transcriptional start site (≥ 15% increase, p ≤ 0.05) and substantially decreased gene expression (fold decrease ≤ 1.5, p ≤ 0.05) in CF eyes compared to SF eyes that have also been linked to human eye disorders. **f-q,** Assessment of *opn1lw1*(f-i), *gnb3a* (j-m), and *crx* (n-q) promoter DNA methylation (f,h,j,l,n,p) and gene expression (g,i,k,m,o,q) in 54 hpf surface (f,g,j,k,n,o) or cave (h,i,l,m,p,q) morphs of *A. mexicanus.* Panels show pie chart graphical representation of the percentage methylation of the promoter CpG (f,h,j,l,n,p), and whole mount *in situ* hybridization of the larval heads (g,i,k,m,o,q) using the probes noted (ventral views, rostral up).

Previous studies have shown that promoter methylation is highly correlated with gene repression^21^. One hundred and twenty-eight genes show substantially increased methylation within the 2 Kb of genomic DNA upstream from the transcriptional start site in CF versus SF (≥ 15% increase, p ≤ 0.05) and decreased expression in CF eyes compared to SF eyes (fold decrease ≤ 1.5, p ≤ 0.05) **(Supplementary data 2)**. These include 39 and 26 genes annotated as having eye expression in mice and humans respectively **(Supplementary data 3)**. Interestingly, nineteen of these genes have been previous linked to human eye disorders **(Fig. 2e)**. These include *opn1lw1*, an opsin associated with cone-rod dystrophy and colorblindness^22,23^, *gnb3a,* defective in autosomal recessive congenital stationary night blindness^24,25^, and *crx*, a photoreceptor-specific transcription factor whose loss leads to blindness in humans^26,27^. Targeted bisulfite sequencing of DNA amplified from the *opn1lw1, gnb3a,* and *crx* promoter regions from 54 hpf SF or CF eyes confirms increased methylation of CpGs in the 5’ upstream sequences of each of these three genes **(Fig. 2f,h,j,l,n,p, Extended Data Fig. 3)**. Whole mount *in situ* hybridization using *opn1lw1, gnb3a,* and *crx* probes also verifies strongly decreased expression of the three genes in CF versus SF eyes **(Fig. 2g,i,k,m,o,q)**. Together, these data show that expression of key human eye disease-associated eye development genes is reduced in Pachón CF compared to their SF relatives, and that many of these eye genes also display increased promoter methylation. Interestingly, the *crx* transcription factor is itself necessary for proper expression of a large number of additional eye genes^27^, many of which are also strongly reduced in CF eyes, even though most of these genes are not themselves methylated **(Extended Data Fig. 4)**.

Increased expression of eye genes such as *opn1lw1, gnb3a,* and *crx* **(Fig. 1l)** and increased eye size **(Fig. 1m)** in *dnmt3bb.1* mutant zebrafish is consistent with the hypothesis that DNA methylation represses eye gene expression to restrain or limit eye development. To experimentally test this idea, we examined whether increasing DNA methylation levels results in decreased eye gene expression and reduced eye development, using previously described zebrafish *Ten-eleven translocation methylcytosine dioxygenase (TET)* mutants that display global DNA hypermethylation^28^. Unlike DNMTs, which add methyl groups to cytosines to generate 5-methylcytosine, TET proteins oxidize methylcytosine to hydroxyl-methylcytosine, promoting DNA demethylation^29,30^. Like mammals, zebrafish have three TET enzymes with partially redundant functions, Tet1, Tet2, and Tet3. Tet2 and Tet3 are the main TET proteins responsible for converting methylcytosine into hydroxymethylcytosine; zebrafish *tet1*^*-/-*^, *tet2*^*-/-*^, *tet3*^*-/-*^ triple mutants do not show additional phenotypes compared to *tet2*^*-/-*^, *tet3*^*-/-*^ double mutants^28^. We found that *tet2*^*-/-*^, *tet3*^*-/-*^ double mutants have smaller eyes than their wild type siblings **(Fig. 3a-c)**. Targeted bisulfite sequencing and qRT-PCR on nucleic acids from whole eyes dissected from 48 hpf *tet2*^*-/-*^, *tet3*^*-/-*^ double mutants and their wild type siblings also revealed increased *crx* and *gnb3a* promoter methylation **(Fig. 3d-g, Extended Data Fig. 5)** and reduced *crx* and *gnb3a* gene expression **(Fig. 3h)** in the double mutants.

**Figure 3.**
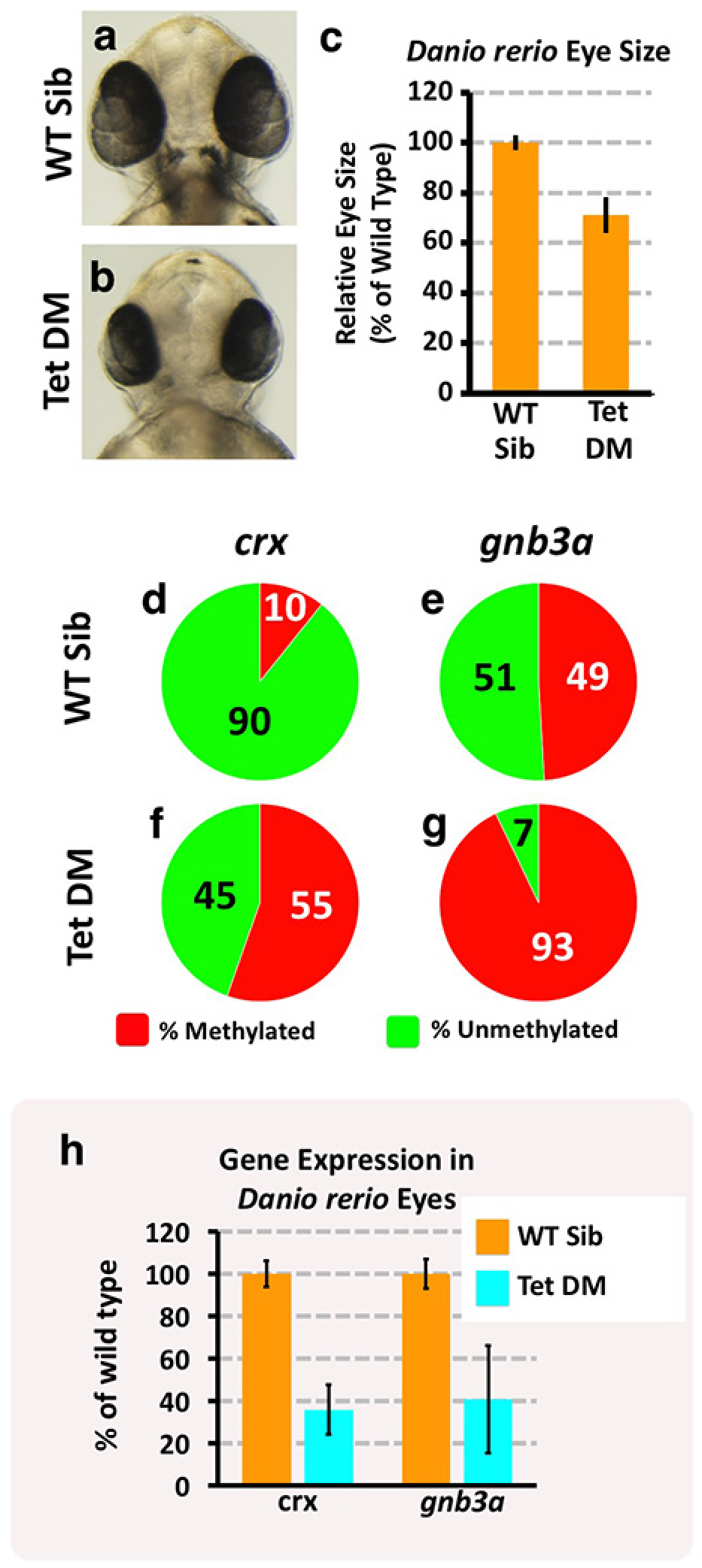
Eye phenotype and associated gene expression changes in wild type and DNA methylation-deficient *Danio rerio*. **a-b,** Transmitted light photomicrographs of the heads of 48 hpf wild type sibling (a) and *tet2*^*-/-*^, *tet3*^*-/-*^ double mutant (b) embryos. **c,** Quantitation of eye size in 48 hpf wild type sibling and *tet2*^*-/-*^, *tet3*^*-/-*^ double mutant embryos. **d-g,** Assessment of *crx* (d,f) and *gnb3a* (e,g), promoter DNA methylation in isolated eyes from 48 hpf wild type sibling (d,e) and *tet2*^*-/-*^, *tet3*^*-/-*^ double mutant (f,g) animals. **h,** Quantitative RT-PCR analysis of the percent relative expression of *crx* and *gnb3a* in isolated eyes from 48 hpf wild type sibling (orange columns) and *tet2*^*-/-*^, *tet3*^*-/-*^ double mutant (blue columns) animals.

To further test whether excess DNA methylation contributes to failure to maintain eye development in Pachón CF, we examined whether eye loss could be “rescued” by pharmacological inhibition of DNA methylation. 5-Azacytidine (AZA) is a well-characterized inhibitor of DNA methylation^31^ approved for treatment of aberrant DNA methylation in myelodysplastic syndrome patients^32^. Since systemic administration of AZA to zebrafish embryos leads to severe pleiotropic defects and early lethality^33^, we carried out single injections of either AZA or control DMSO carrier into the vitreous chamber of the left eye of 42-48 hpf CF embryos, and then scored phenotypes in both injected left and control right eyes at 5 days post fertilization **(Fig. 4a)**. A single early injection of AZA into the left eye resulted in larger eyes than either the control uninjected right eyes in the same animals or the DMSO-injected left eyes of other animals **(Fig. 4b)**. Histological analysis confirmed that AZA-injected 5 dpf CF eyes are significantly larger than uninjected or DMSO injected controls, and that they possess a more organized structure including morphologically well defined lens and retinal layers **(Fig. 4c-e)**.

**Figure 4.**
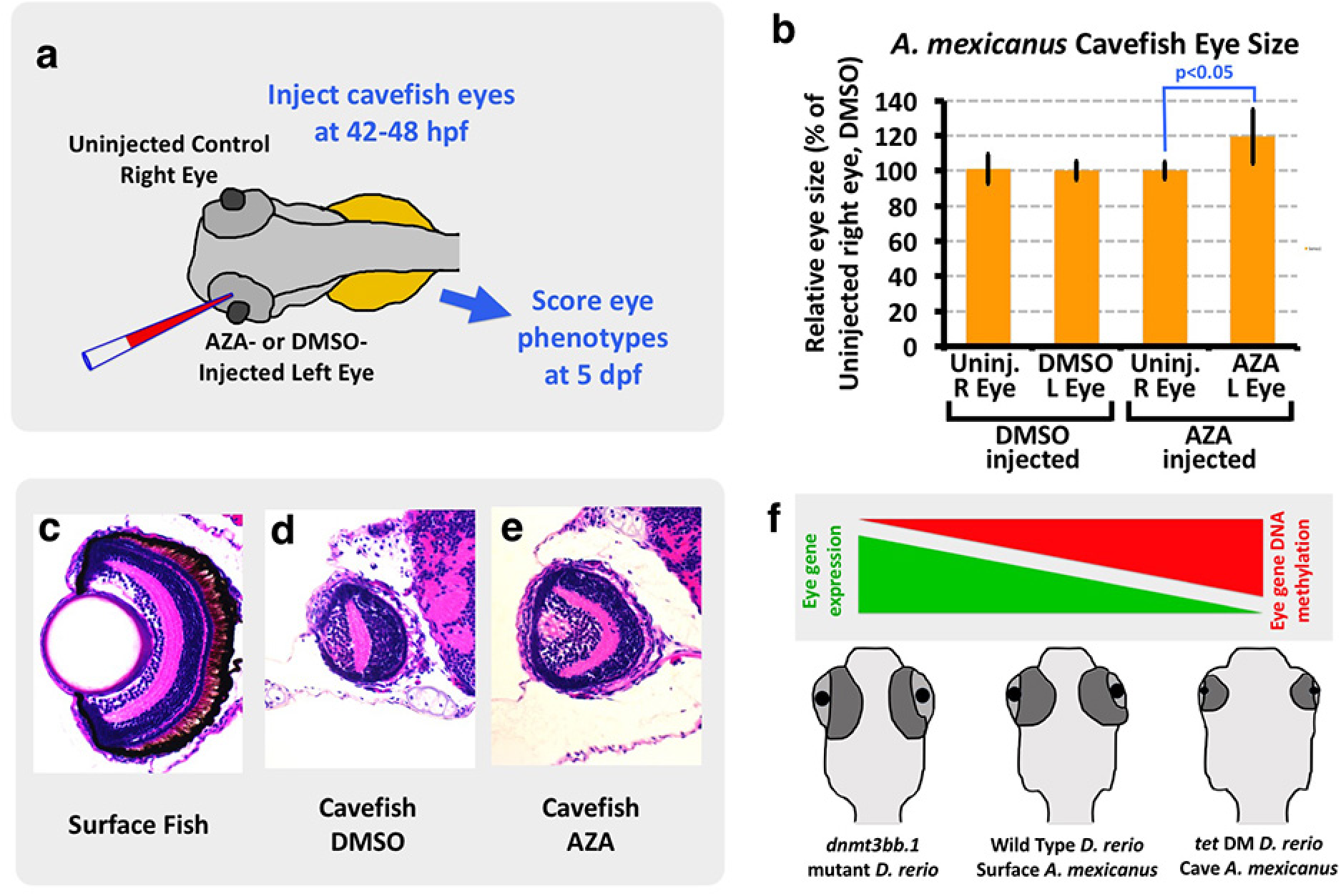
Partial rescue of cavefish eyes by AZA-mediated inhibition of eye DNA methylation. **a**, Schematic diagram showing the experimental procedure for injection of DMSO or AZA into 42-48 hpf cavefish embryo eyes. **b,** Quantitation of eye size in 5 dpf cavefish embryos injected in the left eye with DMSO or AZA. **c-e,** Histological analysis of H&E stained 5 dpf surface fish eye (c), DMSO injected cavefish eye (d) and AZA injected cavefish eye (e). **f,** Model depicting the role of DNA methylation in teleost eye development and degeneration. Hypermethylation and down-regulation of eye gene expression in cavefish and zebrafish *tet*^*2/3*^ double mutants leads to eye degeneration.

Together, our results suggest that DNA methylation plays a critical role in teleost eye development, and that increased DNA methylation-based eye gene repression is a major molecular mechanism underlying CF eye degeneration **(Fig. 4f)**. Many of the key eye genes down-regulated in cave fish are conserved in humans and linked to eye disease and/or blindness, suggesting potential conserved function for these genes across evolution. Although a central role for DNA methylation in development and disease has been well-documented^34,35^, our results suggest that epigenetic processes can play an equally important role in adaptive evolution. Our findings indicate that eye loss in Pachón cavefish occurs via distinct molecular mechanisms compared to naked mole rats, where inactivating mutations are found in multiple key eye genes. This could reflect differences in their evolutionary timescales. Cavefish evolved over the past one to five million years^11^, while naked mole rats evolved seventy-three million years ago^13^, allowing sufficient time to fix and select acquired mutations in genes essential for eye development. It remains to be seen whether epigenetic mechanisms have been used to generate phenotypic variability in other rapidly evolved animals.

## METHODS

Methods, including statements of data availability and any associated accession codes and references, are available in the online version of the paper.

*Note: Any Supplementary Information and Source Data files are available in the online version of the paper.*

## ACKNOWLEDGEMENTS

We thank members of the Weinstein and Jeffery lab for their support, help, and suggestions. We thank NICHD’s Molecular Genomics Laboratory for bisulfite and RNA sequencing assistance. We also thank members of the zebrafish and cavefish community for sharing reagents and protocols. We thank Dr. Karuna Sampath for her comments on the manuscript. We thank Dr. Suzanne McGaugh for her suggestions on cavefish sequence alignments and Dr. Mary Goll for providing the zebrafish *tet2,3* double mutant line. Work in the Weinstein and Jeffery labs is supported by the intramural program of the NICHD and by R01EY024941, respectively.

## AUTHOR CONTRIBUTIONS

A.V.G. and B.M.W. designed the study with inputs from K.A.T. and W.J. A.V.G. and K.A.T. performed the experiments with help from L.M., D.C. and A.D. J.I. analyzed the sequencing data. D.C., A.D., S.S. provided fish husbandry support. A.V.G. and B.M.W. wrote the manuscript with input from all the authors.

**Author Information** RNA-seq data have been deposited in the Gene Expression Omnibus database under accession number XXXXX. Reprints and permissions information is available at www.nature.com/reprints/index.html. The authors declare no competing financial interests. Readers are welcome to comment on the online version of the paper. Correspondence and requests for materials should be addressed to A.V.G. or B.M.W.

## EXTENDED DATA FIGURE LEGENDS

**Extended Data Figure 1.**
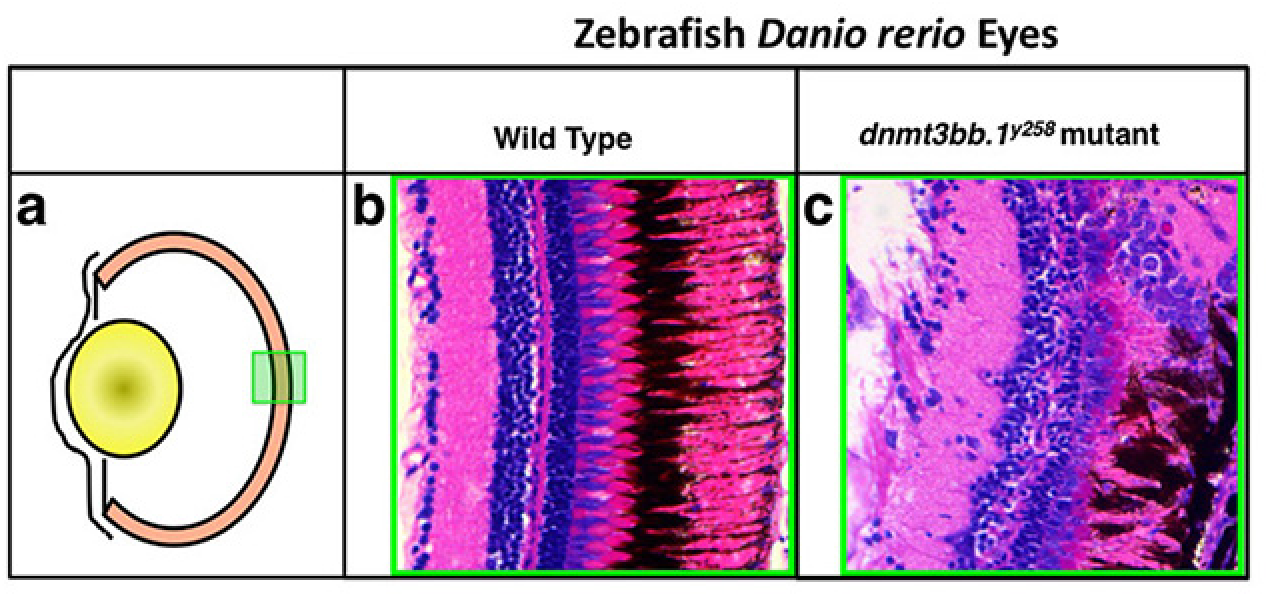
Eye phenotypes in in zebrafish *Danio rerio* wild type and *dnmt3bb.1*^*y258*^ mutant animals. **a**, Schematic diagram of a transverse section through the adult fish eye, with approximate area of the eye shown in panels b and c noted by the green box. **b,c,** H&E-stained transverse sections through adult wild type (b) and *dnmt3bb.1*^*y258*^ mutant (c) eyes. The wild type retina (b) contains well-organized normal layers while hyperplasia and abnormal dysmorphic layers are noted in *dnmt3bb.1*^*y258*^ mutants.

**Extended Data Figure 2.**
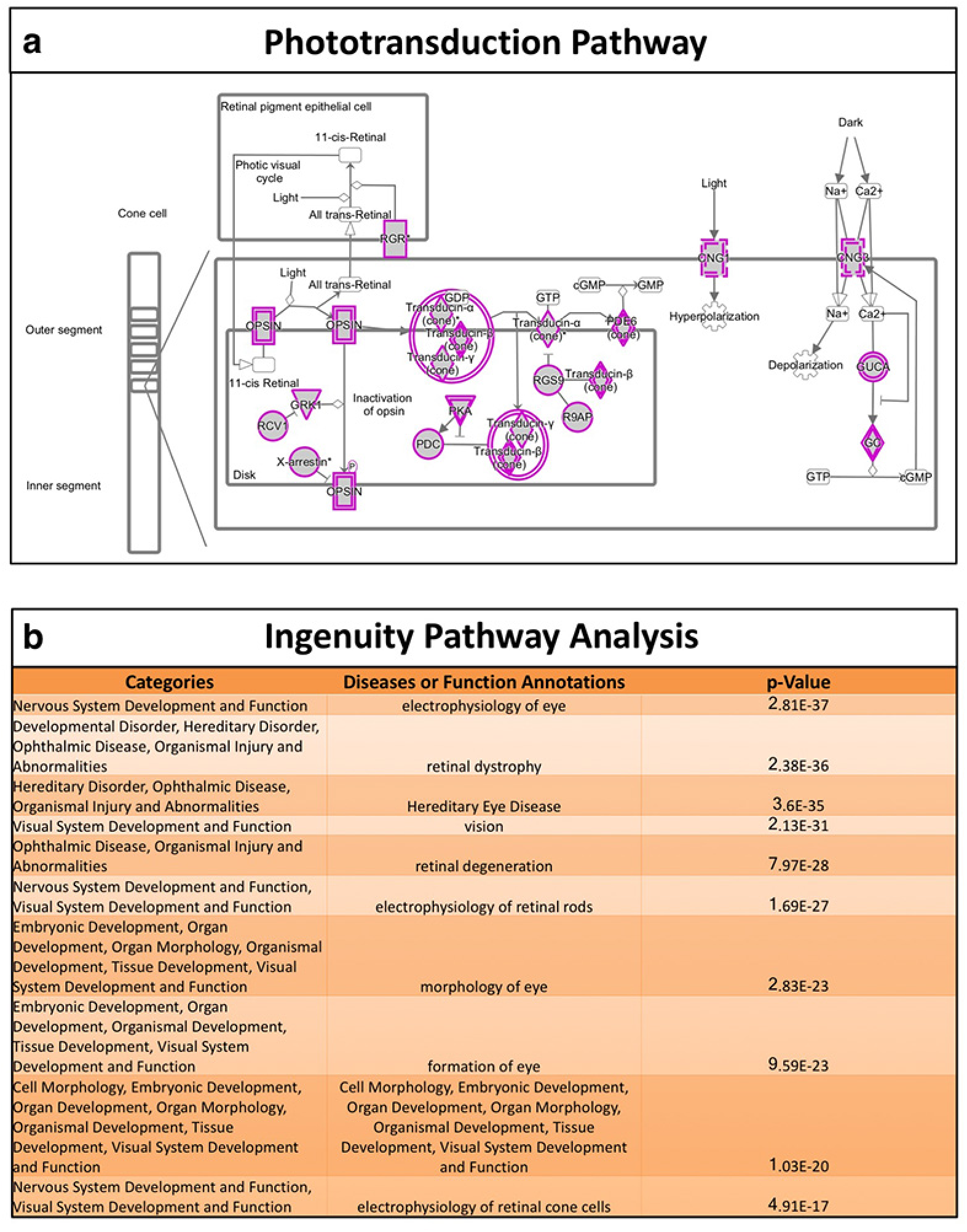
Ingenuity Pathway Analysis (IPA) of differentially expressed genes in cave and surface fish. **a**IPA suggests the phototransduction pathway is one of the key signaling pathways affected in cavefish eyes. Genes highlighted in purple are significantly downregulated. **b,** Developmental processes most significantly affected in cavefish, as predicted by IPA analysis.

**Extended Data Figure 3.**
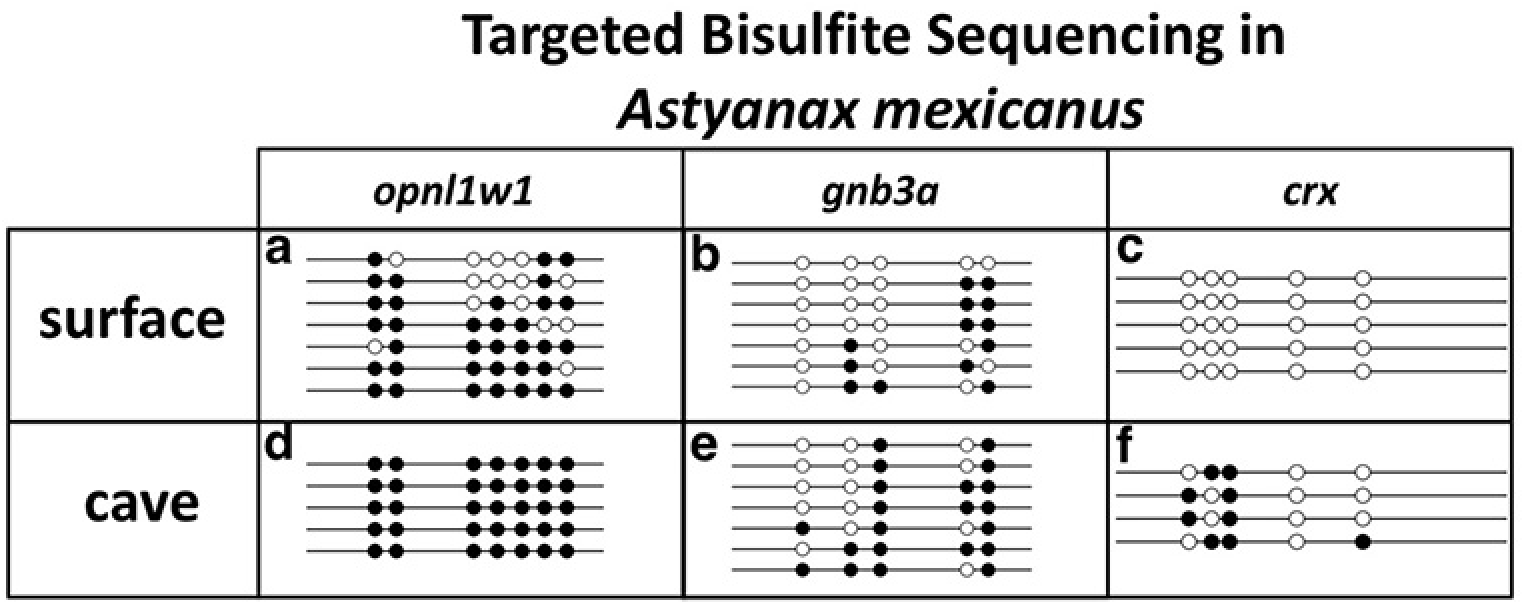
Increased methylation of eye genes in cavefish. **a-f**Targeted bisulfite sequencing analysis of DNA methylation in CpGs isolated from the *opn1lw1* (a,d), *gnb3a* (b,e) and *crx* (c,f) promoter regions of 54-60 hpf surface (a-c) or cavefish (d-f) *Astyanax mexicanus* eyes. Methylation of all three genes is increased in cave compared to surface fish.

**Extended Data Figure 4.**
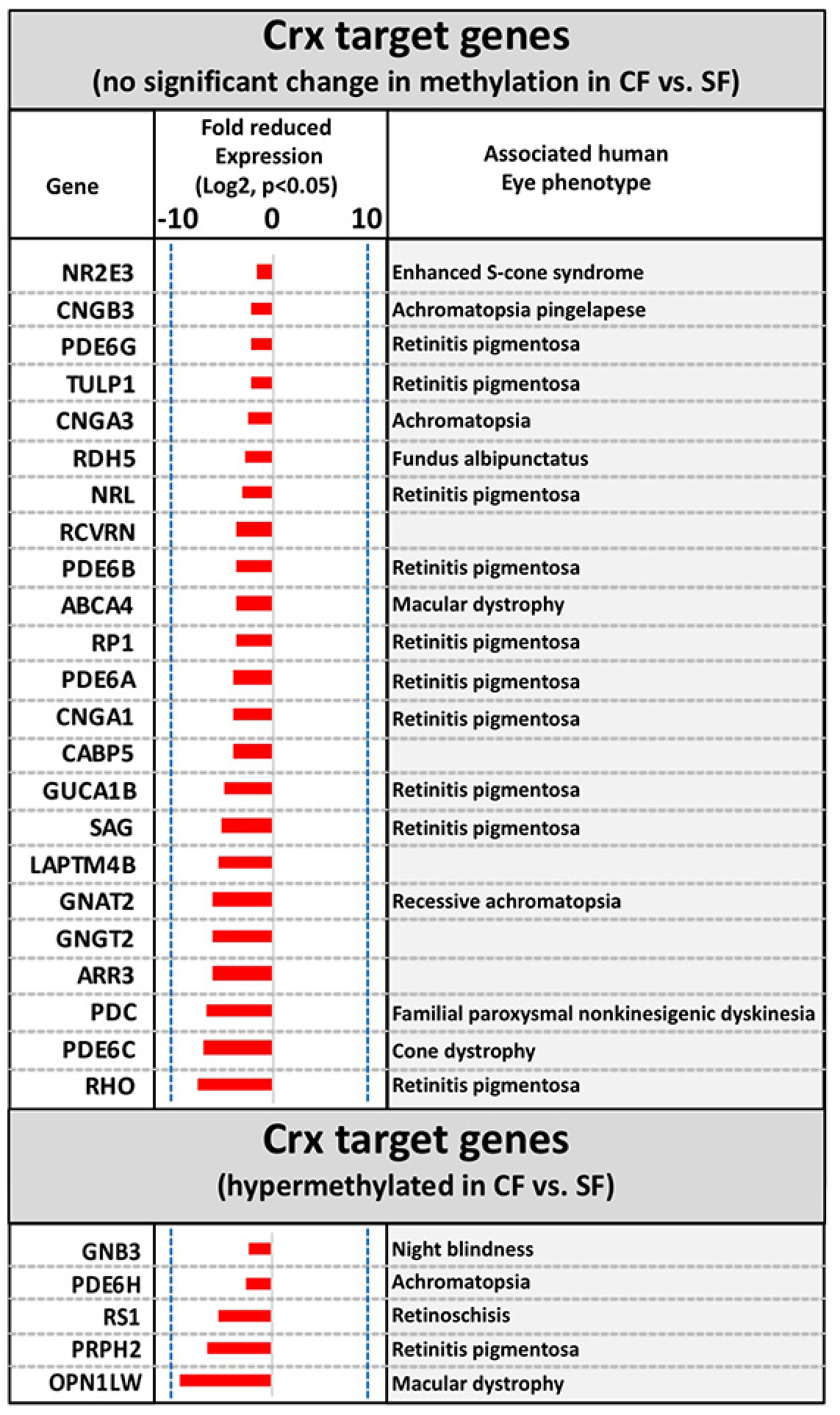
Reduced expression of Crx target genes in cavefish. RNAseq analysis reveals that twenty-eight known Crx target genes also show reduced expression in 54-60 hpf cave versus surface fish eyes (fold decrease ≤ 1.5, p ≤ 0.05), although twenty-three of these twenty-eight genes show no associated changes in DNA methylation. Twenty-three of these genes have also been linked to human eye disorders.

**Extended Data Figure 5.**
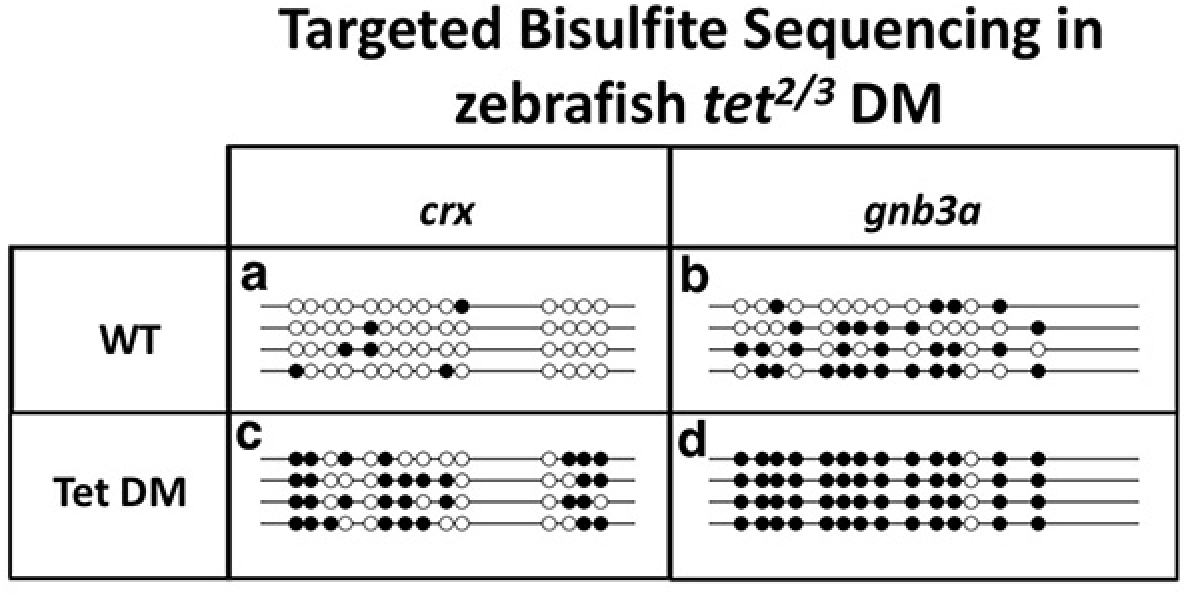
Increased methylation of eye genes in tet2,3 double mutant zebrafish. **a-f**Targeted bisulfite sequencing analysis of DNA methylation in CpG islands isolated from the *crx* (a,c) or *gnb3a* (b,d) promoter regions of 48 hpf wild type sibling (a,b) or *tet2*^*-/-*^, *tet3*^*-/-*^ double mutant (c,d) zebrafish eyes. Methylation of both genes is increased in *tet2*^*-/-*^, *tet3*^*-*^ */-* double mutants compared to their wild type siblings.

## ONLINE METHODS

**Fish stocks and embryos.** Zebrafish lines used in this study include *dnmt3bb.1*^*y258*^ (Ref. 17) and *tet2*^*mk17*^, *tet3*^*mk18*^ (Ref. 28) mutants and the EK wild type line. Surface and Pachon cave populations of *Astyanax mexicanus* are used in this study. Fish were spawned naturally and embryos were raised and staged as described previously^36,37^.

**Genomic DNA isolation, bisulfite conversion, and sequencing.** Surface and cavefish embryos were raised to described stages. Dechorionated embryos were transferred into 1X PBS without calcium and magnesium. Eyes were surgically removed using a pair of fine tip tungsten needles. Total cellular RNA and DNA was isolated from harvested eyes using ZR Duet DNA/RNA miniprep kit (Zymo Research). For whole genome bisulfite sequencing, 200 ng of purified genomic DNA was bisulfite converted using EZ DNA methylation-Lightning kit (Zymo research). Next Gen sequencing libraries were generated from the bisulfite converted DNA using TruSeq DNA methylation kit and TruSeq DNA methylation Index PCR primers (Illumina). Sequencing libraries of two biological replicates from surface and cavefish were run separately on two FlowCells of an Illumina HiSeq2500 sequencer operated in the RapidRun mode with V2 chemistry to yield about 400 million read pair reads (2 X 100bp) for each sample. Raw single-end sequence data was trimmed for quality and adapter sequence using Trimmomatic software, trimming leading or trailing bases at quality < 5 as well as using a sliding window requiring a 4 bp average of quality > 15. Reads trimmed below 50 bp were discarded. Following trimming, reads were aligned using Bismark software against a bisulfite converted Pachon cavefish genome using non-directional alignment. Using defined gene coordinates, gene body and promoter -2kb of TSS regions were assigned and combined conversions and non-conversions were summed to provide an overall bisulfite conversion rate per region. The ratio of conversions was then tested for change between conditions using a two-proportion z-test testing the hypothesis that the proportion of bisulfite conversion had altered between conditions. For targeted bisulfite analysis, bisulfite converted DNA was PCR amplified using primers designed by methprimer^38^ (http://www.urogene.org/cgi-bin/methprimer/methprimer.cgi)One Taq Hotstart 2X master mix in standard buffer DNA polymerase (NEB) was used to amplify bisulfite converted DNA. PCR amplicons were purified and cloned using pCRII-TOPO TA cloning kit (Thermofisher). Miniprep plasmid DNA was sequenced using T7 or SP6 sequencing primers. Sequencing results were analyzed using QUMA^39^ (http://quma.cdb.riken.jp/).

**RNA isolation and sequencing.** Total cellular RNA and DNA was isolated from the harvested eyes using ZR Duet DNA/RNA miniprep kit (Zymo Research). 300-900ng poly-A enriched RNA was converted to indexed sequencing libraries using the TruSeq Standard mRNA library prep kit (Illumina). Libraries for two biological replicates of surface and cavefish were combined and run on one FlowCell of an Illumina HiSeq 2500 sequencer in RapidRun mode with V2 chemistry. 100 million read pairs (2 X 100) per sample were sequenced. Paired-end reads were trimmed using trimmomatic and aligned to Pachon cavefish genome using RNA-STAR, quantitation performed with subread featureCounts and differential expression analysis performed with count data via DESeq2. Human, mouse and zebrafish homologs of cavefish genes were identified using Ensembl BioMart. Ingenuity pathway and Panther GO term enrichment analyses were carried out on RNA sequencing data to identify major signaling pathways and networks affected based on differentially expressed genes.

**RNA isolation, cDNA synthesis and qRT-PCR.** Total cellular RNA was isolated from harvested eyes and other tissues as mentioned in the text using ZR Duet DNA/RNA miniprep kit (Zymo Research). Equal amounts of RNA were converted into cDNA using the ThermoScript RT-PCR system (Invitrogen). Resulting cDNA was used in qPCR using SsoAdvanced™ Universal SYBR^®^ Green Supermix (Biorad) on a CFX96 Real Time system. Primers used in this study are listed in **Supplementary Data 4**.

**Riboprobe synthesis, *In situ* hybridizations and histology.** Antisense riboprobes were generated using Roche DIG and FITC labeling mix. Portions of the coding regions of genes were PCR amplified using One Taq Hotstart 2X master mix in standard buffer DNA polymerase (NEB) and cloned into pCRII-TOPO TA vector (Thermofisher). Sequence verified clones were used to generate antisense riboprobes using appropriate enzymes. The zebrafish *dnmt3bb.1* probe was generated as described previously^17^. Whole mount *in situ* hybridization was carried out as described previously with a few modifications. To reduce non-specific hybridization and enhance signal to noise ratio we used 5% dextran sulfate (Sigma) in the hybridization buffer and pre-adsorbed anti-DIG and anti-FITC antibodies to whole cavefish powder. For histology, embryos and tissue samples were fixed using 4% para-formaldehyde overnight at 4°C and subsequently passed through ascending grades of alcohol followed by paraffin embedding. Sections were stained using hematoxylin and eosin (H&E).

**Microinjection.** Microinjection of 5-Azacytidine (Sigma) into Pachon cavefish embryonic eyes was carried out at 42-48 hpf embryos. Embryos were mounted laterally in low melting point agarose. Injection needles were pulled from filament-containing glass capillaries (World Precision Instruments Cat. No. TW100F-4) using a needle puller (Sutter Instruments). Needles were back loaded and a single 1-2 nl bolus of either 100μM 5-Azacytidine in 5% DMSO or 5% DMSO carrier alone was delivered into the vitreous of the left eye using a Pneumatic Picopump (World Precision Instruments). Embryos were removed from the agarose, allowed to continue to develop until 5 dpf, and then scored for eye size and/or used for histological analysis of the eyes. Injected embryos with axis or brain deformities were discarded from the analysis.

